# Genetic factors underlie the association between anxiety, attitudes and performance in mathematics

**DOI:** 10.1101/719393

**Authors:** Margherita Malanchini, Kaili Rimfeld, Zhe Wang, Stephen A. Petrill, Elliot M. Tucker-Dob, Robert Plomin, Yulia Kovas

**Affiliations:** Department of Biological and Experimental Psychology, Queen Mary University of London; MRC Social, Genetic and Developmental Psychiatry Centre, King’s College London, United Kingdom; Department of Human Development and Family studies, Texas Tech University, United States; Department of Psychology, Ohio State University, Columbus, OH, United States; Department of Psychology, The University of Texas at Austin, United States; Department of Psychology, Goldsmiths, University of London, United Kingdom; International Centre for Research in Human Development, Tomsk State University, Russia

**Author notes:** Corresponding author: Margherita Malanchini, Department of Biological and Experimental Psychology, Queen Mary University of London, Office 2.02, G.E. Fogg Building, Mile End Road, London E1 4NS.

**Keywords:** mathematics anxiety, self-efficacy, mathematics motivation, mathematics cognition, number sense

## Abstract

Students struggling with mathematics anxiety (MA) tend to show lower levels of mathematics self-efficacy and interest as well as lower performance. The current study addresses: (1) how MA relates to different aspects of mathematics attitudes (self-efficacy and interest), ability (understanding numbers, problem-solving ability, and approximate number sense) and achievement (exam scores); (2) to what extent these observed relations are explained by overlapping genetic and environmental factors; and (3) the role of general anxiety in accounting for these associations. The sample comprised 3,410 twin pairs aged 16-21 years, from the Twins Early Development Study. Negative associations of comparable strength emerged between MA and the two measures of mathematics attitudes, phenotypically (∼ -.45) and genetically (∼ -.70). Moderate negative phenotypic (∼ -.35) and strong genetic (∼ -.70) associations were observed between MA and measures of mathematics performance. The only exception was approximate number sense whose phenotypic (-.10) and genetic (-.31) relation with MA was weaker. Multivariate quantitative genetic analyses indicated that all mathematics related measures combined accounted for ∼75% of the genetic variance in MA and ∼20% of its environmental variance. Genetic effects were largely shared across all measures of mathematics anxiety, attitudes, abilities and achievement, with the exception of approximate number sense. This genetic overlap was not accounted for by general anxiety. These results have important implications for future genetic research concerned with identifying the genetic underpinnings of individual variation in mathematics-related traits, as well as for developmental research into how children select and modify their mathematics-related experiences partly based on their genetic predispositions.

## Introduction

Mathematics anxiety (MA) has been consistently linked to lower levels of engagement and motivation and poorer performance in mathematics (1,2). MA is a widespread phenomenon: a recent large-scale investigation of 15-year-olds found that 30% of students across multiple countries part of the Organization for Economic Cooperation and Development (OECD) reported feeling anxious or incapable when solving a mathematics problem (3). Due to the high incidence and hindering consequences for mathematics learning outcomes and experiences (4), it is important to understand the etiology of the association between MA and the attitudinal and performance components of learning mathematics.

The current study investigates the extent to which overlapping genetic and environmental factors underlie the associations between MA, attitudes towards mathematics, cognition and achievement. This work provides a foundation for the search of genetic variants linked to individual differences in MA, and mathematical learning. This study can also inform developmental research into how students select and modify their mathematics-related experiences, partly depending on their genetic predispositions. Moreover, identifying which aspects of performance and attitudes, if any, are more closely associated with anxiety, and the etiologies of these associations, will likely inform the focus of future interventions aimed at reducing MA and fostering mathematics learning.

### Mathematics anxiety and attitudes towards mathematics: self-efficacy and interest

Research has indicated a moderate negative association between MA and mathematics motivation and attitudes, including lower perseverance to learn and practice mathematics (5). Moderate to strong negative associations between mathematics attitudes and anxiety are observed in student populations, as well as in samples of pre-service teachers-trainees working towards obtaining mathematics teaching qualifications-cross culturally (6). The tendency to avoid situations involving mathematics, which covaries with MA is in line with observations of avoidance behavior associated with general anxiety (5,7), and might be related to holding negative beliefs about competence in the subject (8,9). In line with this hypothesis, research found that mathematics self-efficacy, which describes individuals’ perception of their own competency (10), mediated the negative association between MA and performance in high school students (11). Students who achieved higher grades at the start of high school developed higher mathematics self-efficacy, which resulted in lower levels of MA at a two-year follow-up (11). Additionally, self-efficacy was found to mediate the negative association between self-reported self-regulatory behavior and MA: A greater capacity for self-regulation was positively associated with self-efficacy which was in turn negatively linked to MA in an adolescent sample (12).

The expectancy-value theory of motivated behavior (13) proposes that, as well as beliefs and expectancies, subjective task value is a crucial construct characterizing motivated behavior. However, few studies have focused on investigating the association between MA and other aspects of attitudes towards mathematics, beyond self-efficacy. One study found that MA relates to a similar degree to self-efficacy, interest, and importance attributed to mathematics (-.41, -.33, -.30) (14). Similar results were obtained by two previous studies that examined the associaiton bewteen MA and mathemtics importnace, interst and usefulness (15) and bewteen MA and mathematics confidence, interest and importance in a sample of young children (16). Yet, it remains unclear whether the same or distinct genetic and environmental influences underly the relations between MA and mathematics attitudes, such as self-efficacy and interst. The first goal of the present study is to address this gap in the literature.

### Mathematics anxiety and achievement

Students experience MA across the entire distribution of mathematics ability (17). A recent investigation found that, although children with developmental dyscalculia were more likely to show high levels of MA than neurotypical controls, 77% of children presenting with high levels of MA showed average or high performance in mathematics (18). Nevertheless, research has found that students experiencing higher levels of MA on average show weaker mathematical performance. This negative association between MA and mathematics achievement remains significant and moderate after accounting for variation in general cognitive ability (7). The association between MA and achievement has been observed at several developmental stages, from as early as primary school (19,20).

Longitudinal research in an adolescent sample has suggested that the stability of MA increases from moderate to strong during the course of adolescence (21). This observed increase in the stability of MA is partly explained by stable levels of low achievement in mathematics, as achievement was found to drive the development of subsequent MA (21). These results point to the role of negative performance feedback in reinforcing the development of increasingly pervasive levels of MA in adolescence. However, another longitudinal study found reciprocal longitudinal links between negative emotions, including MA, and achievement in mathematics in a sample of secondary school students (22). This is in line with the observation of reciprocal longitudinal links between MA and performance in a sample of primary school students, although effect sizes were observed to be greater for the link from earlier achievement to subsequent anxiety (23). A further longitudinal investigation explored the emergence of the association between mathematics anxiety and achievement in a primary school sample (24). The study found that while no direct longitudinal links between MA and achievement emerged, both constructs were associated with mathematics self-evaluation, suggesting a potential role of attitudes towards mathematics, and particularly self-efficacy, in the development of the link between MA and mathematics achievement (24).

A further line of investigation has explored the possibility that a deficit in lower-level numerical processing may be related to MA via its negative association with mathematics achievement (25). Supporting this hypothesis, two studies have found that high levels of MA were associated with deficits in areas of basic numerical processing such as counting (25) and a simple visual enumeration (26). On the other hand, another investigation (27) failed to find an association between MA and basic numerosity – the ability to discriminate between symbolic and non-symbolic numerical quantities at a first glance (28). Using latent profile analysis, Hart et al. clustered students into different groups, based on their profile in MA, achievement and numerosity. They found that the link between MA and numerosity was weak across all identified groups (27).

Despite the large number of studies on the phenotypic association between MA and mathematics cognition, at present, only one study has explored the association between MA and performance applying a genetically informative design (29). This investigation, conducted in a twin sample, found that the association between MA and performance (measured as mathematics problem-solving ability) was mostly explained by common genetic influences. The second goal of the current study is to extend this research to explore the genetic and environmental overlap between MA and aspects of mathematics attitudes and performance.

### Associations between Mathematics and General Anxiety

Mathematics anxiety and general anxiety partly overlap in their physiological manifestations, which include increased heartbeat, rapid pulse and nervous stomach (30), as well as in cognitive and brain networks (3,9,31). However, the two anxieties are only moderately correlated (.35) (5), suggesting that they may be separate constructs. This is consistent with a number of studies that have observed an association between MA and performance beyond general anxiety (32–34). In line with this, a recent twin study has shown that the genetic and environmental etiology of MA only partly overlap with that of general anxiety (35). Crucially, Wang et al. reported that the partial etiological overlap between MA and general anxiety was independent from the etiology of the overlap between MA and mathematics performance in a problem verification task (29). The third aim of the current study is to examine the extent to which individual differences in general anxiety account for the links between MA and mathematics attitudes, cognition and achievement.

## Methods

### Participants

Participants were members of the Twins Early Development Study (TEDS), a longitudinal study of twins born in the United Kingdom between 1994 and 1996. The families in TEDS are representative of the British population in their socio-economic distribution, ethnicity and parental occupation. Informed consent was obtained from the twins prior to each collection wave. See Haworth et al. (36) for additional information on the TEDS sample. The TEDS study received ethical approval from the King’s College London Ethics Committee. The present study focuses on data collected in a subsample of TEDS twins over two waves: age 16 and age 18-21.

At **age 16**, TEDS twins contributed data on mathematics ability and achievement (*N* = 3,410 pairs, 6,820 twins; MZ = 2,612; DZ = 4,508; 56% females) and mathematics self-efficacy and interest (*N* = 2,505 pairs, 5,010 twins; MZ = 1,954; DZ = 3,270; 61.2% females). At **age 18-21**, the twins contributed data on mathematics anxiety and general anxiety (*N* = 1,506 pairs, 3,012 twins; MZ = 1,172; DZ = 1,846; 63.9% females). All individuals with major medical, genetic or neurodevelopmental disorders were excluded from the dataset.

### Measures

#### Mathematics Anxiety

A modified version of the Abbreviated Math Anxiety Scale (AMAS) (37) was administered to assess mathematics anxiety. The AMAS asks participants to rate how anxious they would feel when facing several mathematics-related situations. The measure includes nine items that are rated on a 5-point scale, ranging from ‘not nervous at all’ to ‘very nervous’. Two items were adapted from the original version to make them age appropriate for the current sample (35), these are: ‘*Listening to a math’s lecture*’ and ‘*Reading a math’s book*’. The AMAS showed excellent internal validity (*α* = .94) and test-retest reliability (*r* = .85) (37).

#### Mathematics attitudes: self-efficacy and interest

Two scales, adapted from the OECD Program for International Student Assessment, measure mathematics self-efficacy and interest. The **mathematics self-efficacy** scale asked participants: ‘*How confident do you feel about having to do the following mathematics tasks?*’ The scale included eight items that participants had to rate on a 4-point scale, from 0 = not at all confident to 3 = very confident. Examples of items are: ‘*Understanding graphs presented in newspapers’*, and ‘*Solving an equation like 3x + 5 = 17’*. The scale showed good internal validity (*α* = .90). The **mathematics interest** scale included three items that participants had to rate on a 4-point scale, from 1 = strongly disagree to 4 = strongly agree. The items were: ‘*I look forward to my mathematics lessons*’; ‘*I do mathematics because I enjoy it*’; and ‘*I am interested in the things I learn in mathematics*’. The scale showed good internal validity (*α* = .93).

#### Mathematics performance

The General Certificate of Secondary Education (GCSE) grades provided a measure of **mathematics exam grade**. The GCSE exams are taken nationwide at the end of the compulsory education, usually when students are 16-years-old. As mathematics is one of the core subjects in the UK educational curriculum, taking the mathematics GCSE exam is a compulsory requirement for all students. Mathematics GCSE scores were collected by questionnaires sent to the twins or their parents by post, via email, or through a phone interview. The GCSE grades, which are given in letters from A* (similar to A+) to G, were re-coded on a scale from 11, corresponding to the highest grade (A*) to 4 corresponding the lowest pass grade (G). No information about ungraded or unclassified results was available. However, these constitute a small proportion of all pupils in the UK (e.g. 1.5% of all exams in 2017; https://www.jcq.org.uk/examination-results/gcses/2017/gcse-full-course-results-summer-2017) and therefore unlikely to constitute a bias in the current study. For 7,367 twins, self- and parent-reported GCSE results were verified using data obtained from the National Pupil database (NPD; www.gov.uk/government/uploads/system/uploads/attachment_data/file/251184/SFR40_2013_FINALv2.pdf), yielding correlations of 0.98 for English, 0.99 for mathematics, and >0.95 for all sciences between self- and parent-reported grades and exam results obtained from NPD (38).

An online test battery assessed mathematics performance with three tests: understanding numbers, problem verification and approximate number sense.

The **understanding numbers** test (39) was developed to specifically assess the ability to understand and solve problems which included numbers and was based on the NFER-Nelson Mathematics 5-14 Series, closely linked to the curriculum requirements in the UK. The items included in the measure were taken from the National Foundation for Education Research (NFER) booklets 8 to 14. The test asked participants to solve 18 mathematics problems arranged in ascending level of difficulty. Questions were presented in multiple formats, ranging from equations to problems. Participants were asked either to type a numerical response into a box or to select one or multiple correct responses out of a set of possible options. An example of one of the difficult items is ‘*Denise has thought of two numbers. The numbers added together make 23. The smaller number subtracted from twice the larger number makes 22. What are Denise’s numbers?*’ with numbers 8 and 15 being correct. Each correct answer was allocated 1 point, resulting in a maximum score of 18. The test showed good reliability in the present sample (α = 0.90).

The **problem verification test** (PVT) (40) presented participants with a series of mathematics equations appearing for 10 seconds on a computer screen. Participants responded to each equation (correct, incorrect, don’t know), by pressing the corresponding keys on the computer keyboard. If they timed out, they were automatically redirected to the following equation. The PVT included 48 items. Examples of items are *‘32 – 16 = 14’*; and *‘2/6 = 3/9’*. Each correct response was allocated the score of 1 and other responses and non-responses the score of 0, for a maximum score of 48. The test showed good reliability in the current sample (α = 0.85).

The approximate **number sense** test (28) included 150 trials displaying arrays of yellow and blue dots, varying in size. Each trial lasted 400 ms and included a different number of blue and yellow dots presented on the screen. Participants were required to judge whether there were more yellow or blue dots on the screen for each trial (see Tosto et al., 2014 for additional information on this task) (41). Each correct answer was allocated the score of 1 and the final score was calculated as the number of correct trials. The final accuracy score correlated strongly (*r* = -.931, p< .0001) with the alternative score calculated using the Weber fraction (42) for which a smaller score indicates better performance.

#### General Anxiety

The Generalized Anxiety Disorder Scale (GAD-7) (43) assessed general anxiety. The scale includes 7 items asking participants to rate on a scale from 1 = not at all to 4 = nearly every day ‘*How often in the past month have you been bothered by the following problems?*’ Examples of items are ‘*Not being able to control worrying’*, and ‘*Feeling afraid as if something awful might happen*’. As well as measuring generalized anxiety disorder, the GAD-7 has been validated and is considered a reliable measure of anxiety in the general population. The GAD-7 is characterized by good internal validity (α = .89) and test-retest reliability *r* = .64 (43).

### Analyses

#### Phenotypic Analyses

Descriptive statistics and ANOVAs were conducted on data from one randomly selected twin out of each pair in order to control for sample dependency (i.e. the fact that the children in the study were twins). Measures were residualized for age and sex and standardized prior to analyses.

#### Genetic Analyses – The Twin Method

The twin method allows for the decomposition of individual differences in a trait into genetic and environmental sources of variance by capitalizing on the genetic relatedness between monozygotic twins (MZ), who share 100% of their genetic makeup, and dizygotic twins (DZ), who share on average 50% of the genes that differ between individuals. The method is further grounded in the assumption that both types of twins who are raised in the same family share their rearing environments to approximately the same extent (44). Comparing how similar MZ and DZ twins are for a given trait (intraclass correlations), it is possible to estimate the relative contribution of genes and environments to variation in that trait. Heritability, the amount of variance in a trait that can be attributed to genetic variance (A), is intuitively calculated as double the difference between the MZ and DZ twin intraclass correlations (45). The ACE model further partitions the variance into shared environment (C), which describes the extent to which twins raised in the same family resemble each other beyond their shared genetic variance, and non-shared environment (E), which describes environmental variance that does not contribute to similarities between twin pairs.

An alternative to the ACE model is the ADE model, which partitions the variance into additive genetic (A), non-additive (or dominant) genetic (D) and non-shared environmental (E) effects. This model is fitted in cases when intraclass correlations for DZ twins are below 50% of the MZ intraclass correlation – indicating non additive genetic influences (46). While additive genetic factors (A) are the sum of the effects of all alleles at all loci contributing to the variation in a trait or to the co-variation between traits, non-additive genetic effects (D) describe interactions between alleles at the same locus (dominance) and at different loci (epistasis). The classic twin design, comparing MZ and DZ twins does not allow to estimate all four sources of influence (A, D, C and E) within one univariate model, as it only includes two coefficients of relatedness (47). Therefore, with the classic twin design it is possible to partition the variance into three sources of influences: A, E, and either C or D.

ACE models were fitted for mathematics GCSE, understanding numbers, and mathematics problems verification test. For these measures, intraclass correlations for DZ pairs were more than half of those for MZ pairs, suggesting that environmental factors contributed to the similarity between twins beyond their genetic similarity.

ADE models were fitted for MA, general anxiety, mathematics interest, mathematics self-efficacy, and number sense. For these measures, the DZ intraclass correlation were less than half that of MZ, indicating non additive genetic effects.

The twin method can be extended to the exploration of the covariance between two or more traits (***multivariate genetic analysis***). Multivariate genetic analysis allows for the decomposition of the covariance between multiple traits into genetic and environmental sources of variance, by modelling the cross-twin cross-trait covariances. Cross-twin cross-trait covariances describe the association between two variables, with twin 1 score on variable 1 correlated with twin 2 score on variable 2, which are calculated separately for MZ and DZ twins.

One way of partitioning the genetic and environmental covariation between two or more traits is to conduct a ***multivariate Cholesky decomposition***. The Cholesky decomposition allows to examine the overlapping and independent genetic (A), shared (C) (or non-additive D), and non-shared (E) environmental effects on the variance in two or more traits (48). A Cholesky decomposition can be interpreted similarly to a hierarchical regression analysis, as the independent contribution of a predictor variable to the dependent variable is estimated after accounting for the variance it shares with other predictors previously entered in the model. The current study applies Cholesky decompositions to the investigation of the genetic and environmental overlap between MA, mathematics motivation and performance.

## Results

### Descriptive statistics and sex differences

Descriptive statistics for all variables, which were normally distributed, are presented in the supplementary Table S1. Due to previously reported sex differences in mathematics anxiety (49) and performance (50), we firstly tested for sex differences in all measures using univariate ANOVAs (Table S2). Males showed significantly higher levels of mathematics self-efficacy, interest and performance across all measures, and lower levels of mathematics and general anxiety. Sex explained a relatively small portion of the variance in all measures (0-7%). Previous genetically informative work on these same measures (35,51) did not find support for the existence of qualitative differences in the etiology of mathematics anxiety and performance between males and females. Consequently, these analyses were not repeated. Table S3 reports the twin correlations separately for same-sex (DZss) and opposite-sex (Dzos) dizygotic pairs. As can be seen in Table S3, the twin correlations for DZss and Dzos are similar for all variables, suggesting no qualitative or quantitative sex differences in ACE estimates.

### Genetic and environmental variation in mathematics related traits

Eight univariate models were conducted in order to partition individual differences in all mathematics-related traits. Figure 1 reports the results of these univariate analyses. (Table S4 reports intraclass correlations and 95% confidence intervals for these univariate analyses.) With the exception of GCSE exam scores (for which significant C was found), the AE model was found to be the best fit for the data for all traits. Dropping the C or D paths did not significantly decrease the goodness of fit of the univariate models (see Table S5). Estimates of heritability ranged between 36% and 63%, and the remaining variance was explained by non-shared environmental factors, which also include measurement error. Shared environmental factors accounted for 18% of the variance in GCSE exam scores.

**Figure 1.**
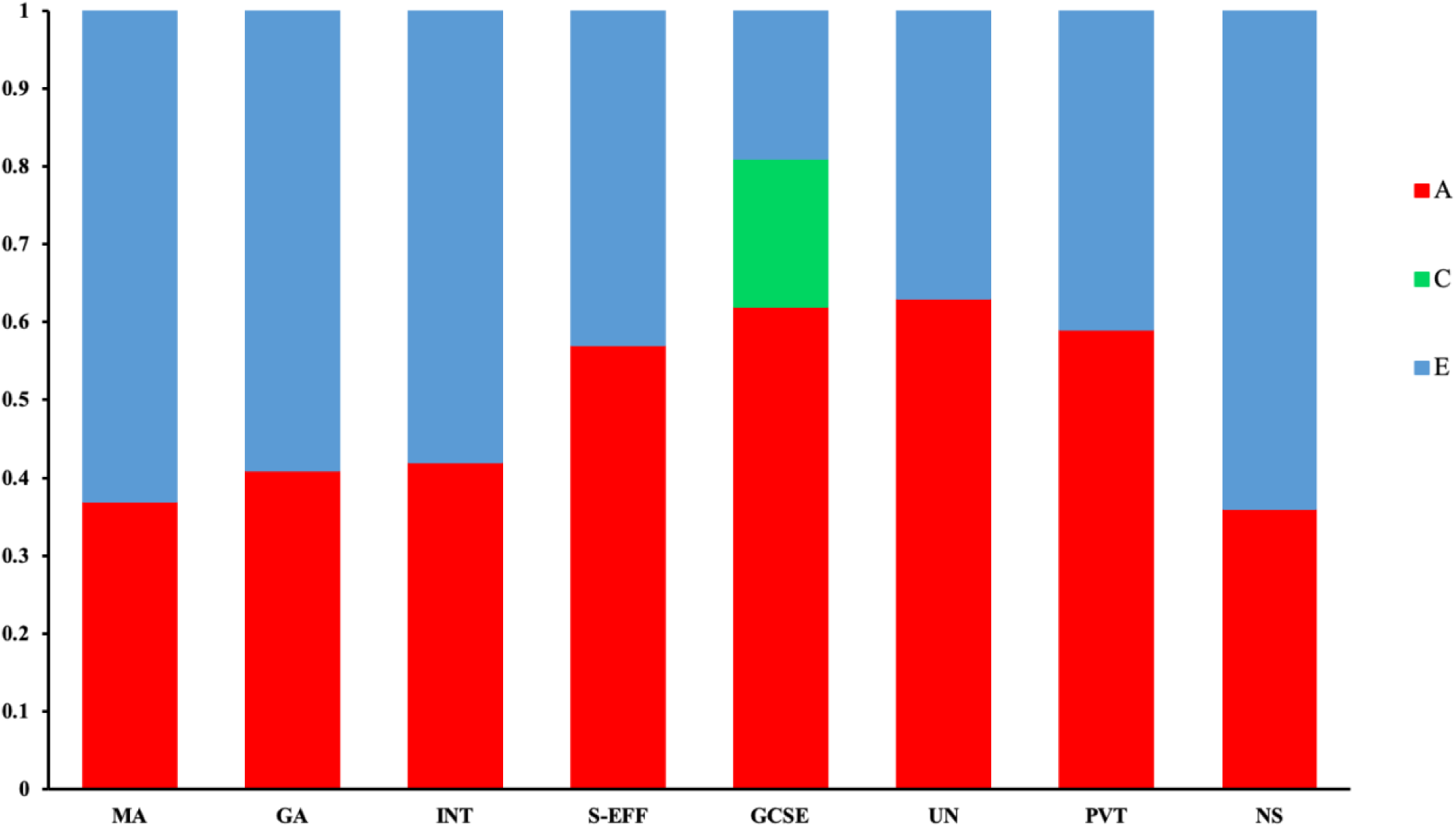
Univariate genetic (A), shared environmental (C) and non-shared environmental (E) estimates for all mathematics related measures; MA = mathematics anxiety; GA = general anxiety; INT = interest; S-EFF = self-efficacy; GCSE = mathematics GCSE exam score; UN = understanding numbers; PVT = problem verification test; NS = number sense.

### Phenotypic and genetic correlations across all mathematics related traits

Figure 2 presents the phenotypic (observed) and genetic correlations between all mathematics related traits. Moderate negative phenotypic correlations (*r* ranging between -.31 and -.45) and strong negative genetic correlations (*rA* ranging between -.67 and -.75) were observed between MA and all other mathematics related variables. The only exception was the association between MA and approximate number sense, which was weak phenotypically (*r* = -.10) and modest genetically (*rA* = -.31). Measures of mathematics attitudes shared a positive moderate to strong phenotypic association (*r* ranging between .38 and .56) and strong genetic association (*rA* ranging between .57 and .82) with measures of mathematics performance. Phenotypic and genetic correlations across measures of mathematics performance were strong. Again, an exception was approximate number sense, which was only moderately related to other mathematical measures.

**Figure 2.**
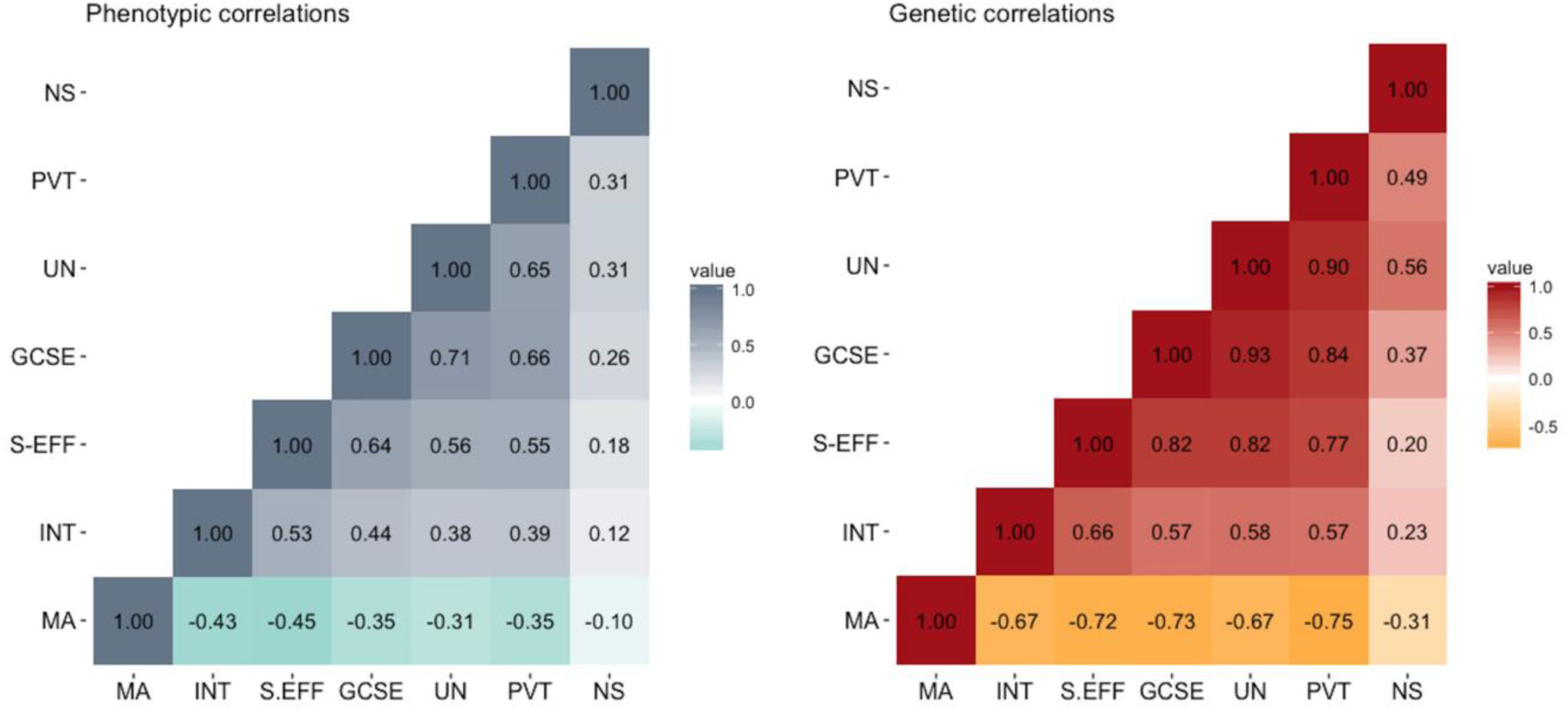
Phenotypic and genetic correlations across all mathematics related measures; MA = mathematics anxiety; INT = interest; S-EFF = self-efficacy; GCSE = mathematics GCSE exam score; UN = understanding numbers; PVT = problem verification test; NS = number sense.

### Multivariate associations between MA and mathematics attitudes

We conducted two Cholesky decomposition analyses to explore the unique genetic and environmental overlap between each measure of mathematics self-efficacy and interest and MA. Following the rationale of hierarchical regression, in order to explore the unique genetic and environmental variance shared between self-efficacy and MA after accounting for interest, we entered interest as the first variable in the model, followed by self-efficacy and MA (Figure 3a). The attitudes variables were then inverted in a second model (Figure 3b), which explored the unique association between interest and MA after accounting for self-efficacy. Both models found that MA shared ∼35% of its genetic variance with mathematics attitudes, and these shared genetic effects were common across both measures of mathematics attitudes. The percentage of genetic variance in MA that overlaps with self-efficacy and interest can be calculated dividing the effect size of the standardize a1,3 paths in Figure 3a and 3b (.13) by the total heritability of MA (.37). Specific genetic associations between each attitude construct and MA were smaller in magnitude, accounting for between 5% and 8% of additional genetic variance in MA.

**Figure 3.**
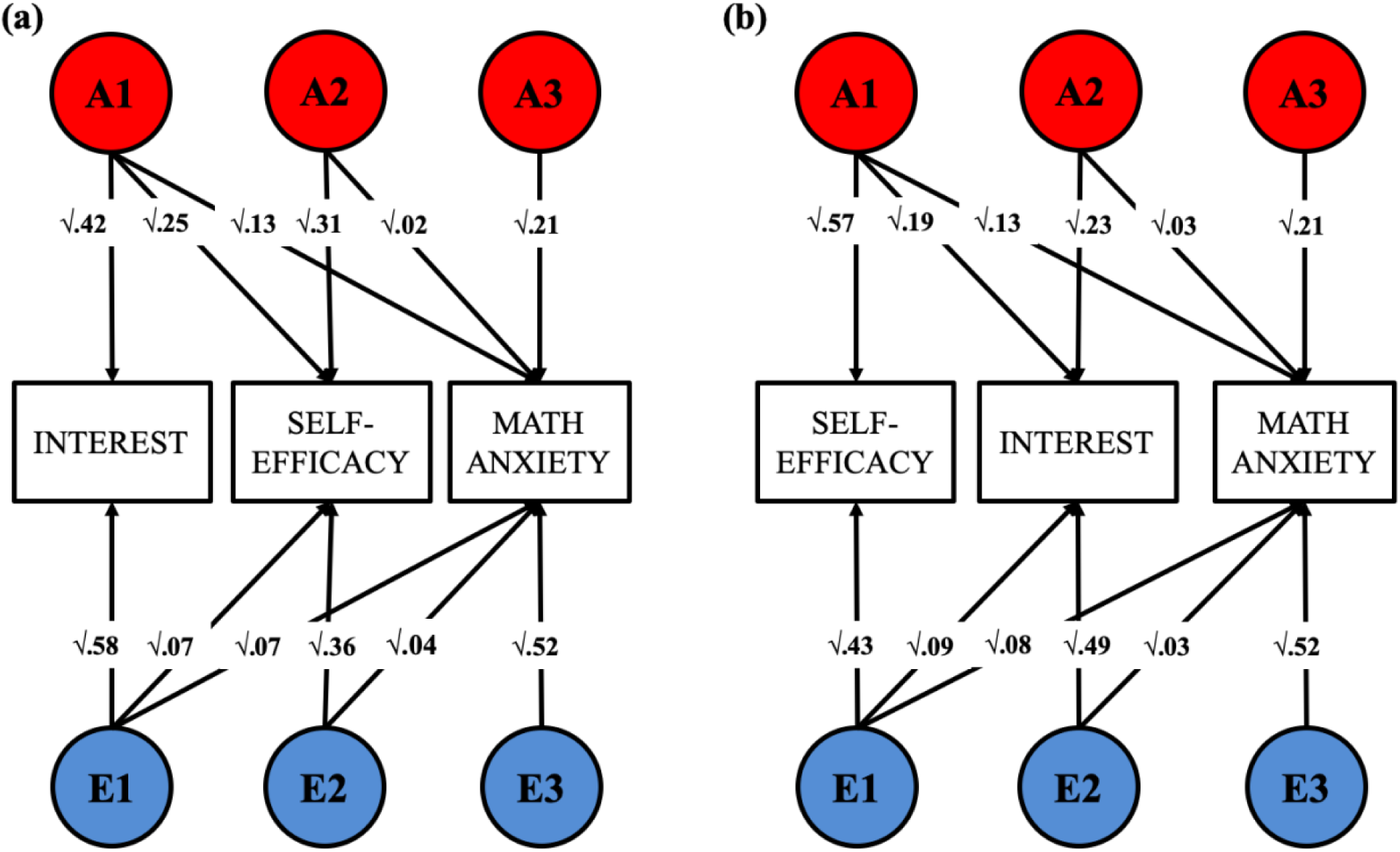
Trivariate Cholesky decompositions exploring the unique genetic and non-shared environmental overlap between MA and mathematics self-efficacy (**2a**), and MA and mathematics interest (**2b**) - after accounting for the other measure of motivation, entered at the first stage in the model.

A degree of specificity was observed in the non-shared environmental overlap between the measures, as ∼10% of the non-shared environmental variance in MA overlapped with self-efficacy independently of the other ∼10% of the variance it shared with interest. These can be calculated diving the effect size of the standardized e1,2 and e1,3 paths linking self-efficacy and interest to MA (.09 and .08, respectively) by the total non-shared environmental variance in MA (.63).

### Genetic and environmental variance common to MA, mathematics attitudes and performance

Two additional Cholesky decompositions explored association between mathematics anxiety and all mathematics-related traits. The first decomposition (Figure 4) explored the genetic and environmental variance that MA shared with each of the other mathematics related measure – with MA entered first in the analysis. Results of this first decomposition indicated that the heritability of MA accounted for between 35% and 50% of the genetic variance in the mathematics related measures, with the exception of approximate number sense, for which only 8% of the genetic variance overlapped with MA. The weak shared environmental variance, which could not be dropped from this multivariate analysis (see Table S6) was shared with other mathematics-related traits. In contrast, the non-shared environmental variance in MA accounted for a small proportion of non-shared environmental variance in all other mathematics related measures (between 0 and 10%). Full results for this multivariate model are reported in Table S7.

**Figure 4.**
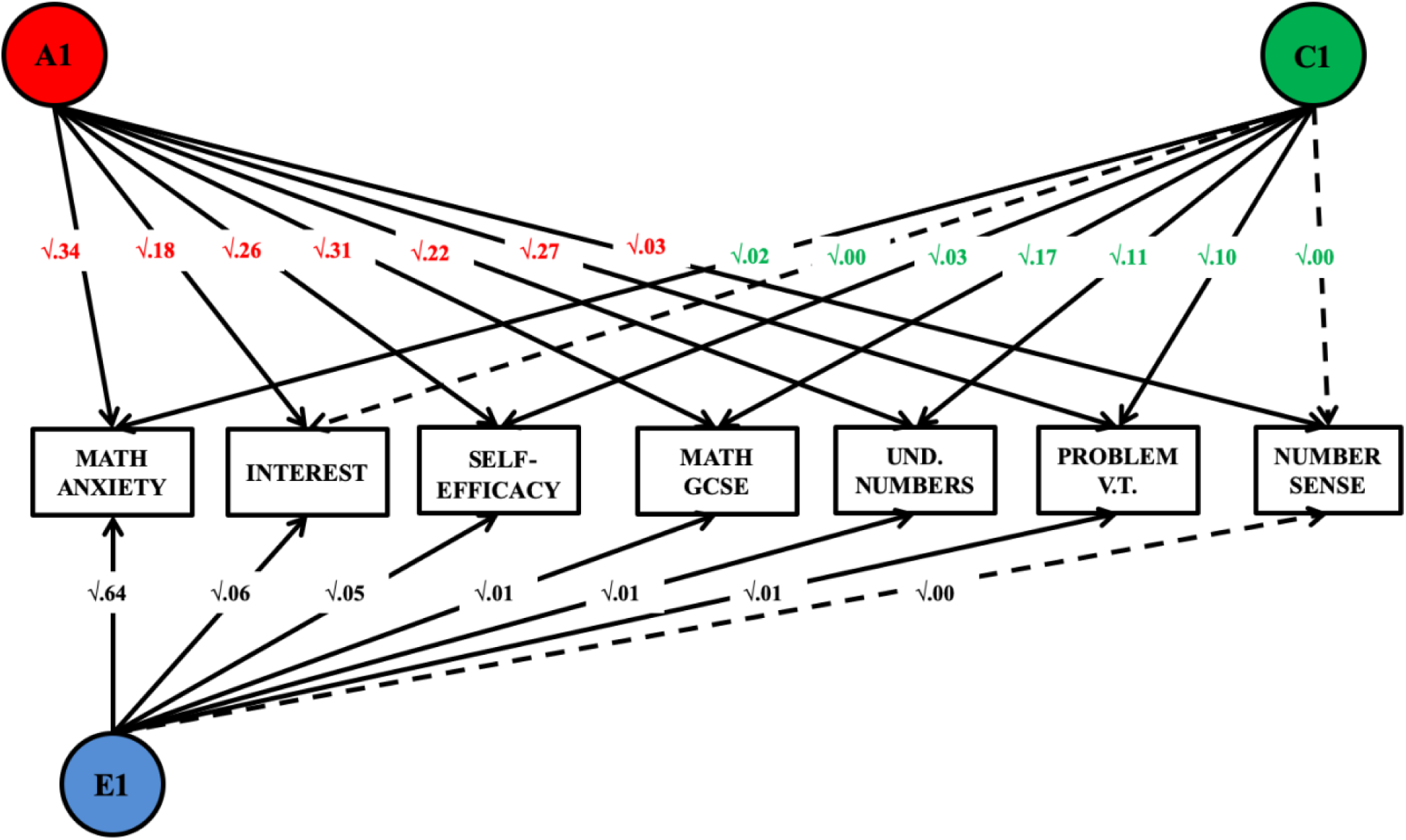
Proportion of genetic and environmental variance shared between MA and all other measures of mathematics motivation and performance. For ease of reading and interpretation, the current figure shows only the A1 genetic paths and C1 and E1 environmental paths. These standardized and squared path estimates were derived from a full Cholesky decomposition (see results Table S7).

The second multivariate model (Figure 5) included the same seven variables but entered in a different order - providing a different perspective on examining the genetic and environmental overlap between MA, attitudes and performance in mathematics. This second analysis tested how much of the genetic and environmental variance in MA was accounted for by all the other variables previously entered in the model, and how much remained specific to MA. In order to explore whether there was specificity in the genetic and environmental variance shared between measures of mathematics affect after accounting for performance, all measures of mathematics performance were entered first in the model, followed by measures of attitudes and, lastly, MA. The results (see Table S8 for the results of the full Cholesky decomposition) indicated that 76% of the genetic variance in MA was shared with the other mathematics related measures, and that the majority of this substantial genetic overlap was common to measures of performance and attitudes. The two mathematics attitudes measures accounted for an additional 10% of this shared genetic variance. The small shared environmental variance in MA was entirely shared with mathematics performance (GCSE exam scores). In contrast, most (83%) of the non-shared environmental variance was specific to MA, with 8% in common with measures of mathematics performance and 9% in common with measures of attitudes.

**Figure 5.**
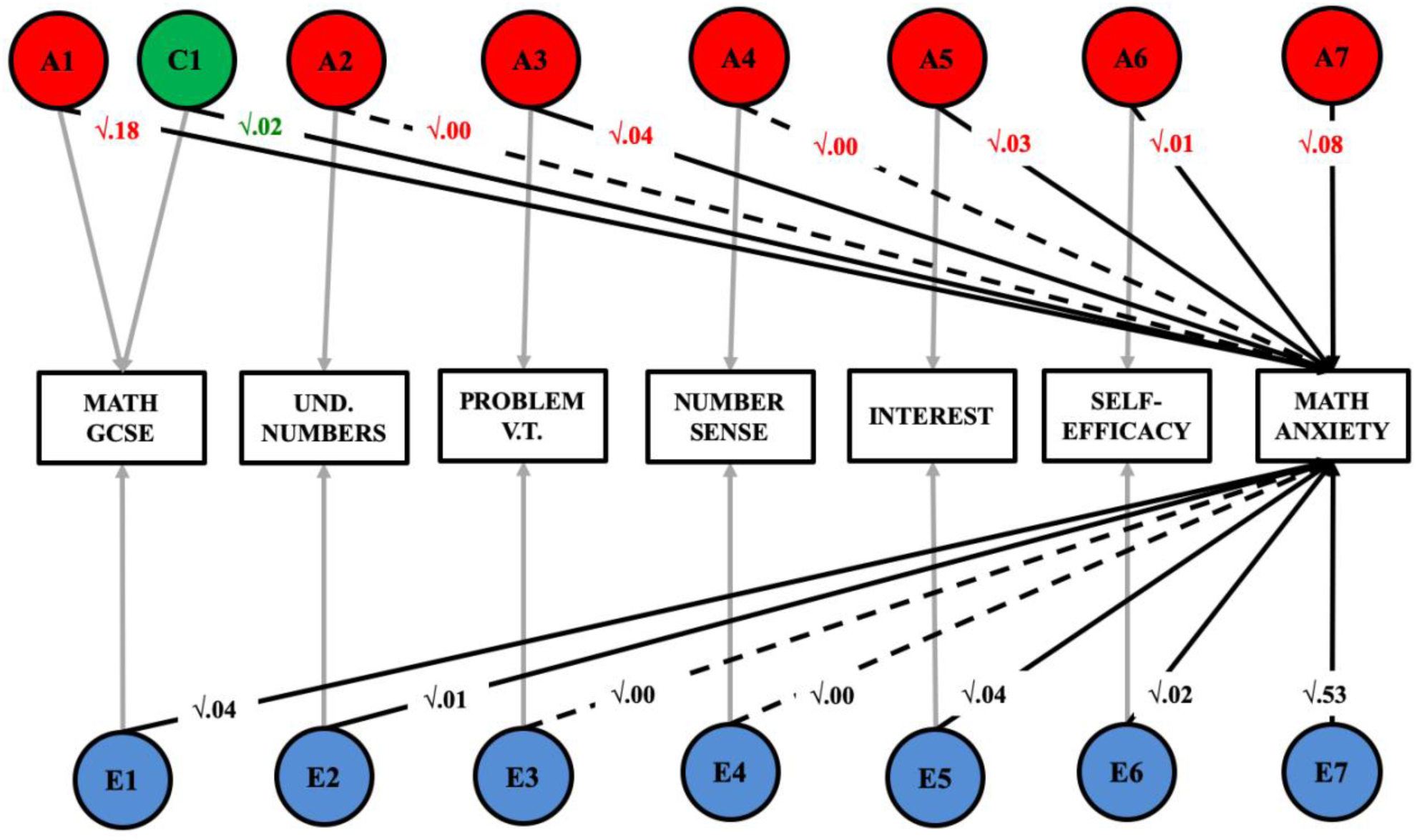
Proportion of genetic and environmental variance in MA accounted for by all other mathematics-related measures. For ease of reading and interpretation, the current figure shows only the standardized and squared path estimates linking each predictor to variation in MA - derived from a full Cholesky decomposition (see full results Table S8). The results of this decomposition can be interpreted as those of a hierarchical regression: the effect of each predictor is estimated after accounting for the variance explained by each other predictor previously entered in the model.

### The role of general anxiety in the MA-attitudes-performance association

The Cholesky decomposition presented in Figure 4 was repeated including general anxiety, in order to test whether the observed genetic and environmental associations between MA and mathematics attitudes and performance could be accounted for by general anxiety. Results (Table S9) indicated that general anxiety accounted for 22% of the genetic variance and 4% of the environmental variance in MA. However, after accounting for general anxiety, the genetic and environmental associations between MA, attitudes and performance remained mostly unchanged.

## Discussion

The present investigation was the first to adopt a genetically informative framework to explore the genetic and environmental overlap between anxiety, self-efficacy, interest and performance in the domain of mathematics, and the role of general anxiety in accounting for the observed associations. The results showed a substantial genetic overlap between all mathematics related traits. This shared genetic variance was largely independent from general anxiety.

The first aim of the study was to explore whether MA was similarly associated with different measures of mathematics attitudes. Results indicated that similar effect size characterized the negative associations between MA and mathematics self-efficacy and interest. More than one third of the genetic variance in MA overlapped with mathematics self-efficacy and interest. In contrast, environmental effects across MA, and attitudes towards mathematics were mostly specific to each measure.

These results show that a high degree of generality characterizes the genetic overlap between mathematics anxiety, interest and self-efficacy, as largely overlapping genetic effects were found to contribute to variation in all constructs. On the other hand, the environmental links between mathematics anxiety and interest and self-efficacy were found to be largely specific to each construct, and to include individual-specific, or stochastic processes (including measurement error), which are encompassed by non-shared environmental variance, rather than family-wide characteristics which are subsumed under shared environmental effects. In fact, the majority of the non-shared environmental links were specific to the pairwise associations between MA and self-efficacy and MA and interest, and not shared across the three constructs.

Different environmental experiences, such as different classrooms, teachers, peers, life events, or perception of parental involvement and socio-economic status, could all play a role in explaining these observed non-shared environmental associations. Evidence of an overlap between environmental factors across measures of mathematics attitudes and anxiety is consistent with research showing that the classroom learning environment is similarly associated with both MA and mathematics self-efficacy (52). Future research is needed to identify the environmental factors that link MA to self-efficacy but not interest, and vice versa.

The second aim of the study was to explore the common genetic and environmental variance across MA and multiple measures of mathematics attitudes and performance. Common genetic factors were observed to characterize the etiology of all mathematics-related traits. MA accounted for more than one third of the genetic variance in mathematics attitudes and between one third and half of the genetic variance in mathematics performance. In turn, measures of mathematics performance accounted for three quarters of the genetic variance in MA. These differences in the proportion of heritability accounted for by MA and mathematics performance likely reflect the difference in heritability between the measures. MA, as it is often observed for self-reported constructs (53,54), is moderately heritable, while the heritability of mathematics performance is more substantial. Longitudinal studies in genetically informative samples (e.g. 51) are needed to investigate causal directions between constructs.

A significant genetic association between mathematics attitudes, particularly self-efficacy, and performance beyond MA was observed. The only exception was approximate number sense, which shared a very small proportion of its genetic variance with all measures of mathematics affect. The negligible association between approximate number sense and MA is in line with previous evidence (27). Moreover, our findings suggest that MA is particularly linked to numerical tasks that involve learned symbolic representations of discrete quantitates, rather than approximate representations (41). The lack of a shared genetic etiology between measures of mathematics affect and approximate number sense suggests that basic approximate numerical skills are unlikely to function as a cognitive precursor of the negative association observed between MA and performance.

The third aim of the present study was to explore whether the association between mathematics anxiety, attitudes and performance was domain specific, or whether general anxiety accounted for part of their association. Although general anxiety and MA shared ∼20% of their genetic variance, general anxiety did not account for the association between MA and measures of mathematics attitudes and performance; in fact, it was mostly unrelated to variation in mathematics interest, self-efficacy and performance. These findings extend the line of evidence provided by Wang et al. (2014) and suggest that the common etiology of the association between MA, self-efficacy, interest and cognition may be partly specific to the domain of mathematics. Our results are consistent with evidence showing genetic and environmental specificity for general anxiety and measures of anxiety in the mathematics and spatial domains (35). Research integrating measures of anxiety and performance in other domains, such as for example second language learning, will be able to further test the hypothesis of domain-specific factors linking affect and cognition in the field of mathematics.

The current investigation presents some limitations. As well as relying on the methodological assumptions of twin design (see Rijsdijk & Sham, 2002 for a detailed description) (47), the models employed in the current investigation do not specifically account for gene–environment interplay. One possibility is that the observed genetic association between MA, attitudes and performance may operate via environmental effects that are correlated or interact with genetic predisposition. For example, children with a genetic predisposition towards experiencing difficulties with mathematics may develop a greater vulnerability to negative social influences in the context of mathematics, such as negative feedback received from teachers or parents on their effort and performance, which in turn may lead to greater feelings of anxiety towards mathematics (56). This has the potential to generate a negative feedback loop (7) between performance, motivation and anxiety - that is potentially a product of interacting inherited and environmental factors. The present investigation, including one time point for each measure of mathematics anxiety, attitudes and performance does not allow us to establish the direction of causality between the observed associations. Longitudinal genetically informative studies, integrating multiple measures of mathematics attitudes, anxiety and performance are therefore needed.

A further limitation of the present investigation is that the measure of MA was not collected at the same time as the measures of mathematics performance and motivation. However, longitudinal investigations found moderate to strong phenotypic and genetic stability of MA (21), attitudes and performance (57), which suggests that the links between this two-year time lapse capture the majority of the processes involved.

## Conclusions

The present investigation was the first to examine the genetic and environmental overlap between MA and several aspects of mathematics attitudes and performance. Our findings of a shared, likely domain-specific, etiology between these mathematics-related traits provide a seminal step for future genetic research aimed at identifying the specific genes implicated in variation in the cognitive and non-cognitive factors of mathematics. Our results suggest that the majority of genetic variants implicated in individual differences across mathematics anxiety, attitudes and performance are unlikely to be implicated in variation in general anxiety. The current findings also provide a starting ground for developmental research to delve deeper into the observed common genetic links, examining how the experiential processes through which children select, shape and modify their mathematical experiences interact with genetic predispositions to produce variation in mathematics anxiety, attitudes and performance.

## Acknowledgements

We gratefully acknowledge the ongoing contribution of the participants in the Twins Early Development Study (TEDS) and their families. TEDS is supported by a program grant to R.P. from the UK Medical Research Council (MR/M021475/1 and previously G0901245), with additional support from the US National Institutes of Health (AG046938). The research leading to these results has also received funding from the European Research Council under the European Union’s Seventh Framework Programme (FP7/2007-2013)/ grant agreement n. 602768. R.P. is supported by a Medical Research Council Professorship award (G19/2). M.M.’s work is partly supported by the David Wechsler Early Career grant for innovative work in cognition and by NIH grant R01HD083613 awarded to E.T.D. Y.K.’s work is supported by the Tomsk State University competitiveness improvement programme. The funders had no role in study design, data collection and analysis, decision to publish, or preparation of the manuscript.

Supplementary material, meant for online publication, accompanies the paper.

## References

1. Devine A, Fawcett K, Szűcs D, Dowker A. Gender differences in mathematics anxiety and the relation to mathematics performance while controlling for test anxiety. Behav Brain Funct. 2012 Jan;8(1):33.

2. Eden C, Heine A, Jacobs AM. Mathematics Anxiety and Its Development in the Course of Formal Schooling—A Review. Psychology [Internet]. 2013;04(06):27–35.

3. Suárez-Pellicioni, Macarena, Núñez-Peña, María Isabel, & Colomé Á. Math anxiety: A review of its cognitive consequences, psychophysiological correlates, and brain bases. Cogn Affect Behav Neurosci. 2016

4. Carey E, Hill F, Devine A, Szücs D. The chicken or the egg? The direction of the relationship between mathematics anxiety and mathematics performance. Front Psychol. 2016;6(JAN):1–6.

5. Hembree R. The Nature, Effects, and Relief of Mathematics Anxiety. J Res Math Educ. 1990;21(1):33–46.

6. Hoffman B. “I think I can, but I’m afraid to try”: The role of self-efficacy beliefs and mathematics anxiety in mathematics problem-solving efficiency. Learn Individ Differ. 2010 Jun;20(3):276–83.

7. Ashcraft MH, Moore AM. Mathematic Anxiety and the Effective Drop in Performance. J Psychoeducxational Assess. 2009;27(3):197–205.

8. Ashcraft MH. Math Anxiety: Personal, Educational, and Cognitive Consequences. Curr Dir Psychol Sci. 2002;11(5):181–5.

9. Ashcraft MH, Kirk EP. The relationships among working memory, math anxiety, and performance. J Exp Psychol Gen. 2001;130(2):224–37.

10. Bandura A. Self-efficacy: Toward a Unifying Theory of Behavioral Change. Psychol Rev Univ [Internet]. 1977;Vol. 84(No. 2):191–215.

11. Meece JL, Wigfield A, Eccles JS. Predictors of math anxiety and its influence on young adolescents’ course enrollment intentions and performance in mathematics. J Educ Psychol. 1990;82(1):60–70.

12. Jain S, Dowson M. Mathematics anxiety as a function of multidimensional self-regulation and selfefficacy. Contemp Educ Psychol. 2009;34(3):240–9.

13. Wigfield A, Eccles JS. Expectancy-Value Theory of Achievamanr Motivation. Contemp Educ Psychol. 2000;25(1):68–81.

14. Wang Z, Shakeshaft N, Schofield K, Malanchini M. Anxiety is not enough to drive me away: A latent profile analysis on math anxiety and math motivation. PLoS One. 2018;13(2).

15. Wigfield A, Meece JL. Math Anxiety in Elementary and Secondary School Students. J Educ Psychol. 1988;80(2):210–6.

16. Ganley CM, McGraw AL. The development and validation of a revised version of the math anxiety scale for young children. Front Psychol. 2016;7(AUG).

17. Ashcraft MH, Krause JA. Working memory, math performance, and math anxiety. Psychon Bull Rev [Internet]. 2007;14(2):243–8.

18. Devine A, Hill F, Carey E, Szűcs D. Cognitive and emotional math problems largely dissociate: Prevalence of developmental dyscalculia and mathematics anxiety. J Educ Psychol. 2018;110(3):431–44.

19. Ramirez G, Gunderson EA, Levine SC, Beilock SL. Math Anxiety, Working Memory, and Math Achievement in Early Elementary School. J Cogn Dev. 2013;14(2):187–202.

20. Vukovic RK, Kieffer MJ, Bailey SP, Harari RR. Mathematics anxiety in young children: Concurrent and longitudinal associations with mathematical performance. Contemp Educ Psychol [Internet]. 2013;38(1):1–10.

21. Ma X, Xu J, Xu J. The causal ordering of mathematics anxiety and mathematics achievement: a longitudinal panel analysis. J Adolesc. 2004 Apr;27(2):165–79.

22. Pekrun R, Lichtenfeld S, Marsh HW, Murayama K, Goetz T. Achievement Emotions and Academic Performance: Longitudinal Models of Reciprocal Effects. Child Dev. 2017;88(5):1653–70.

23. Gunderson EA, Park D, Maloney EA, Beilock SL, Levine SC. Reciprocal relations among motivational frameworks, math anxiety, and math achievement in early elementary school. J Cogn Dev [Internet]. 2018;19(1):21–46.

24. Krinzinger H, Kaufmann L, Willmes K. Math Anxiety and Math Ability in Early Primary School Years. J Psychoeduc Assess [Internet]. 2010;27(3):206–25.

25. Maloney EA, Ansari D, Fugelsang JA. Rapid Communication.The effect of mathematics anxiety on the processing of numerical magnitude. Q J Exp Psychol. 2011;64(1):10–6.

26. Maloney EA, Risko EF, Ansari D, Fugelsang J. Mathematics anxiety affects counting but not subitizing during visual enumeration. Cognition. 2010;114(2):293–7.

27. Hart S, Logan JAR, Thompson LA, Kovas Y, Mcloughlin G, Petrill S. A latent profile analysis of math achievement, numerosity, and math anxiety in twins. Florida State Univ Libr. 2015;Faculty Pu:1–48.

28. Halberda J, Ly R, Wilmer JB, Naiman DQ, Germine L. Number sense across the lifespan as revealed by a massive Internet-based sample. Proc Natl Acad Sci U S A. 2012 Jul;109(28):11116–20.

29. Wang Z, Hart SA, Kovas Y, Lukowski S, Soden B, Thompson L a, et al. Who is afraid of math? Two sources of genetic variance for mathematical anxiety. J Child Psychol Psychiatry. 2014 Mar;1–9.

30. Adams C. Overcoming math anxiety. Math Intell. 2001;23:49–50.

31. Wu SS, Barth M, Amin H, Malcarne V, Menon V. Math anxiety in second and third graders and its relation to mathematics achievement. Front Psychol. 2012;3(June):1–11.

32. Cargnelutti E, Tomasetto C, Passolunghi MC. How is anxiety related to math performance in young students? A longitudinal study of Grade 2 to Grade 3 children. Cogn Emot. 2017;31(4):755–64.

33. Wu SS, Willcutt EG, Escovar E, Menon V. Mathematics Achievement and Anxiety and Their Relation to Internalizing and Externalizing Behaviors. J Learn Disabil. 2014;47(6):503–14.

34. Hill F, Mammarella IC, Devine A, Caviola S, Passolunghi MC, Szucs D. Maths anxiety in primary and secondary school students: Gender differences, developmental changes and anxiety specificity. Learn Individ Differ. 2016;48:45–53.

35. Malanchini M, Rimfeld K, Shakeshaft NG, Rodic M, Schofield K, Selzam S, et al. The genetic and environmental aetiology of spatial, mathematics and general anxiety. Sci Rep. 2017;7(42218).

36. Haworth CMA, Davis OSP, Plomin R. Twins Early Development Study (TEDS): A Genetically Sensitive Investigation of Cognitive and Behavioral Development From Childhood to Young Adulthood. Twin Res Hum Genet [Internet]. 2013;16(01):117–25.

37. Hopko DR, Mahadevan R, Bare RL, Hunt MK. The abbreviated math anxiety scale (AMAS). Assessment. 2003;10(2):178–82.

38. Krapohl E, Rimfeld K, Shakeshaft NG, Trzaskowski M, McMillan A, Pingault J-BJ-B, et al. The high heritability of educational achievement reflects many genetically influenced traits, not just intelligence. Proc Natl Acad Sci [Internet]. 2014;111(42):15273–8.

39. Tosto MG, Asbury K, Mazzocco MMM, Petrill SA, Kovas Y. From classroom environment to mathematics achievement: The mediating role of self-perceived ability and subject interest. Learn Individ Differ [Internet]. 2016;50:260–9.

40. Murphy MM, Mazzocco MMM. Mathematics Learning Disabilities in Girls With Fragile X or Turner … J Learn Disabil. 2008;29–46.

41. Tosto MG, Petrill SA, Halberda J, Trzaskowski M, Tikhomirova TN, Bogdanova OY, et al. Why do we differ in number sense? Evidence from a genetically sensitive investigation. Intelligence. 2014;43(1):35–46.

42. Halberda J, Mazzocco MMM, Feigenson L. Individual differences in non-verbal number acuity correlate with maths achievement. Nature [Internet]. 2008 Sep 7;455:665.

43. Löwe B, Decker O, Müller S, Brähler E, Schellberg D, Herzog W, et al. Validation and Standardization of the Generalized Anxiety Disorder Screener (GAD-7) in the General Population. Med Care [Internet]. 2008;46(3):266–74.

44. Kendler KS, Neale MC, Kessler RC, Heath AC, Eaves L. A twin study of recent life events and difficulties. Arch Gen Psychiatry. 1993;50:789–96.

45. Martin NG, Eaves LJ. Stages; the First To Determine the Genetical and Environmental Model. Most. 1977;38:79–95.

46. Eaves LJ, Silberg JL, Meyer JM, Maes HH, Simonoff E, Pickles A, et al. Genetics and Developmental Psychopathology: 2. The Main Effects of Genes and Environment on Behavioral Problems in the Virginia Twin Study of Adolescent Behavioral Development. J Child Psychol Psychiatry. 1997;38(8):965–80.

47. Rijsdijk F V, Sham PC. Analytic approaches to twin data using structural equation models. Brief Bioinform [Internet]. 2002;3(2):119–33.

48. Neale MC, Boker SM, Bergeman CS, Maes HH. The Utility of Genetically Informative Data in the Study of Development. 2005;1–58.

49. Goetz T, Bieg M, Lüdtke O, Pekrun R, Hall NC. Do Girls Really Experience More Anxiety in Mathematics? Psychol Sci. 2013 Aug.

50. Reilly D, Neumann DL, Andrews G. Sex differences in mathematics and science achievement: A meta-analysis of National Assessment of Educational Progress assessments. J Educ Psychol [Internet]. 2015;107(3):645–62.

51. Tosto MG, Hanscombe KB, Haworth CMA, Davis OSP, Petrill SA, Dale PS, et al. Why do spatial abilities predict mathematical performance? Dev Sci. 2014;17(3):462–70.

52. Taylor BA, Fraser BJ. Relationships between learning environment and mathematics anxiety. Learn Environ Res. 2013;16(2):297–313.

53. Malanchini M, Engelhardt LE, Grotzinger AD, Harden KP, Tucker-drob EM. “Same But Different”: Associations Between Multiple Aspects of Self-Regulation, Cognition and Academic Abilities. J Pers Soc Psychol [Internet]. 2018;advance on.

54. Briley DA, Tucker-Drob EM. Genetic and environmental continuity in personality development: A meta-analysis. Vol. 140, Psychological Bulletin; 2014. p. 1303–31.

55. Malanchini M, Wang Z, Voronin I, Schenker VJ, Plomin R, Petrill SA, et al. Reading self-perceived ability, enjoyment and achievement: A genetically informative study of their reciprocal links over time. Dev Psychol. 2017;53(4).

56. Beilock SL, Maloney E a. Math anxiety: a factor in math achievement not to be ignored. Policy Insights from Behavorial Brain Sci. 2015;2(1):4–12.

57. Luo YLL, Kovas Y, Haworth CMA, Plomin R. The etiology of mathematical self-evaluation and mathematics achievement: Understanding the relationship using a cross-lagged twin study from ages 9 to 12. Learn Individ Differ [Internet]. 2011;21(6):710–8.

