## Supplementary Information for "Genetic factors underlie the association between anxiety, attitudes and performance in mathematics"

**List of Supplementary Tables:**

**Table S1.** Descriptive statistics for all the variables included in the study
**Table S2**. Univariate analyses of variance (ANOVAs) examining sex differences in all variables
**Table S3.** Intra-class correlations and cross-twin cross-trait correlations for same-sex DZ twins (above diagonal) and opposite-sex DZ twins (below diagonal)
**Table S4.** Univariate ACE estimates including 95% confidence intervals
**Table S5**. Model fit indices for all univariate models and nested models
**Table S6**. Model fit indices for the multivariate Cholesky decompositions exploring the associations between mathematics anxiety, attitudes and performance
**Table S7**. Standardised paths for the Cholesky decomposition exploring the origins of the association between mathematics anxiety, mathematics attitudes and mathematics performance.
**Table S8**. Standardised squared paths for the Cholesky decomposition exploring the origins of the association between mathematics performance, attitudes and mathematics anxiety.
**Table S9**. Standardised paths for the Cholesky decomposition exploring the origins of the association between general anxiety, mathematics anxiety, mathematics attitudes and mathematics performance.

**Table S1.** Descriptive statistics for all the variables included in the study

| | MA | GA | INT | S-EFF | GCSE | UN | PVT | NS |
|---|---|---|---|---|---|---|---|---|
| N* | 1457 | 1457 | 2506 | 2505 | 3410 | 2237 | 2345 | 2470 |
| Mean | 2.27 | 1.97 | 2.54 | 17.71 | 8.91 | 11.55 | 36.08 | 115.75 |
| St Dev | 1.00 | 0.74 | 0.94 | 5.47 | 1.46 | 4.33 | 6.69 | 9.75 |
| Skew | 0.79 | 0.84 | -0.07 | -0.90 | -0.55 | -0.77 | -0.58 | -0.64 |
| SE Skew | 0.06 | 0.06 | 0.05 | 0.05 | 0.04 | 0.05 | 0.05 | 0.49 |
| Kurtosis | -0.16 | 0.03 | -0.99 | 0.28 | 0.29 | 0.15 | -0.21 | 0.25 |
| SE Kurt | 0.13 | 0.13 | 0.10 | 0.10 | 0.08 | 0.10 | 0.10 | 0.09 |
| Minimum | 1 | 1 | 1 | 0 | 4 | 0 | 15 | 79 |
| Maximum | 5 | 4 | 4 | 24 | 11 | 18 | 48 | 140 |

Note: St Dev = standard deviation; SE = standard error; * one twin out of each pair was randomly selected; MA = mathematics anxiety; GA= general anxiety; INT = interest; S-EFF = self-efficacy; GCSE = mathematics GCSE exam score; UN = understanding numbers; PVT = problem verification test; NS = number sense

**Table S2**. Univariate analyses of variance (ANOVAs) examining sex differences in all variables

| | *Female M (SD), N* | *Male M (SD), N* | *F* | *Partial η2* |
|---|---|---|---|---|
| GA | 2.09 (.77), *N* = 938 | 1.74 (.62) , *N* = 519 | 77.15** | 0.05 |
| MA | 2.45 (1.04), *N* = 938 | 1.91 (.79) , *N* = 519 | 101.58** | 0.07 |
| INT | 2.42(.95), *N* = 1474 | 2.69(.89) , *N* = 1032 | 48.37** | 0.02 |
| S-EFF | 16.51(5.60), *N* = 1473 | 19.40(4.78) , *N* = 1032 | 181.65** | 0.07 |
| GCSE | 8.80 (1.48), *N* = 1812 | 9.03 (1.41) , *N* = 1598 | 20.48** | 0.06 |
| UN | 11.00 (4.40), *N* = 1317 | 12.35(4.08) , *N* = 920 | 54.13** | 0.02 |
| PVT | 34.81(6.46), *N* = 1364 | 37.85(6.58) , *N* = 981 | 124.700** | 0.05 |
| NS | 115.63(9.72), *N* = 1383 | 115.91(9.85) , *N* = 976 | .455(ns) | 0.00 |

*Note*: one twin out of each pair was selected to control for non-independence of observation; ** = $p < .01$; MA = mathematics anxiety; GA = general anxiety; INT = interest; S-EFF = self-efficacy; GCSE = mathematics GCSE exam score; UN = understanding numbers; PVT = problem verification test; NS = number sense.

**Table S3.** Intra-class correlations and cross-twin cross-trait correlations for same-sex DZ twins (above diagonal) and opposite-sex DZ twins (below diagonal)

| | **MA tw1** | **GA tw1** | **Int. tw1** | **S-Eff tw1** | **GCSE tw1** | **UN tw1** | **PVT tw1** | **NS tw1** | **MA tw2** | **GA tw2** | **Int. tw2** | **S-Eff tw2** | **GCSE tw2** | **UN tw2** | **PVT tw2** | **NS tw2** |
|---|---|---|---|---|---|---|---|---|---|---|---|---|---|---|---|---|
| **MA tw1** | - | 0.29 | -0.36 | -0.40 | -0.36 | -0.24 | -0.28 | -0.11 | 0.09 | 0.09 | -0.10 | -0.09 | -0.08 | -0.04 | -0.07 | -0.08 |
| **GA tw1** | 0.31 | - | -0.02 | 0.00 | -0.02 | 0.03 | 0.01 | 0.00 | 0.09 | 0.25 | -0.01 | -0.03 | 0.00 | 0.02 | -0.05 | 0.00 |
| **Int. tw1** | -0.45 | -0.04 | - | 0.53 | 0.44 | 0.36 | 0.40 | 0.21 | 0.00 | 0.02 | 0.24 | 0.17 | 0.25 | 0.20 | 0.11 | 0.11 |
| **S-Eff tw1** | -0.43 | -0.09 | 0.54 | - | 0.61 | 0.55 | 0.53 | 0.21 | -0.02 | -0.01 | 0.16 | 0.32 | 0.36 | 0.29 | 0.19 | 0.13 |
| **GCSE tw1** | -0.40 | -0.14 | 0.47 | 0.64 | - | 0.69 | 0.64 | 0.27 | 0.02 | 0.04 | 0.11 | 0.32 | 0.57 | 0.39 | 0.29 | 0.21 |
| **UN tw1** | -0.33 | -0.13 | 0.40 | 0.54 | 0.70 | - | 0.65 | 0.30 | 0.04 | 0.07 | 0.05 | 0.27 | 0.39 | 0.40 | 0.29 | 0.19 |
| **PVT tw1** | -0.38 | -0.13 | 0.39 | 0.51 | 0.63 | 0.61 | - | 0.30 | 0.02 | 0.02 | 0.09 | 0.22 | 0.34 | 0.36 | 0.28 | 0.19 |
| **NS tw1** | -0.11 | -0.08 | 0.11 | 0.20 | 0.30 | 0.30 | 0.29 | - | 0.06 | 0.00 | 0.00 | 0.04 | 0.11 | 0.13 | 0.06 | 0.15 |
| **MA tw2** | 0.05 | -0.05 | -0.09 | -0.06 | -0.04 | -0.05 | -0.05 | -0.04 | - | 0.34 | -0.39 | -0.44 | -0.29 | -0.29 | -0.31 | -0.07 |
| **GA tw2** | 0.05 | 0.07 | -0.09 | -0.12 | -0.06 | -0.02 | -0.04 | -0.09 | 0.26 | - | -0.12 | -0.13 | -0.09 | -0.03 | -0.08 | 0.01 |
| **Int. tw2** | -0.05 | -0.03 | 0.12 | 0.15 | 0.16 | 0.11 | 0.14 | 0.08 | -0.43 | -0.10 | - | 0.49 | 0.39 | 0.36 | 0.36 | 0.14 |
| **S-Eff tw2** | -0.07 | -0.04 | 0.10 | 0.17 | 0.22 | 0.18 | 0.16 | 0.13 | -0.44 | -0.15 | 0.57 | - | 0.61 | 0.55 | 0.54 | 0.20 |
| **GCSE tw2** | -0.13 | -0.03 | 0.17 | 0.27 | 0.43 | 0.35 | 0.29 | 0.13 | -0.36 | -0.14 | 0.46 | 0.64 | - | 0.68 | 0.63 | 0.28 |

| | | | | | | | | | | | | | | | | |
|---|---|---|---|---|---|---|---|---|---|---|---|---|---|---|---|---|
| **UN tw2** | -0.07 | -0.01 | 0.08 | 0.15 | 0.25 | 0.28 | 0.21 | 0.12 | -0.32 | -0.08 | 0.34 | 0.55 | 0.67 | - | 0.64 | 0.32 |
| **PVT tw2** | -0.07 | -0.05 | 0.11 | 0.19 | 0.29 | 0.32 | 0.28 | 0.08 | -0.40 | -0.09 | 0.40 | 0.49 | 0.63 | 0.64 | - | 0.35 |
| **NS tw2** | -0.02 | -0.14 | 0.08 | 0.08 | 0.11 | 0.13 | 0.10 | 0.22 | -0.08 | -0.12 | 0.08 | 0.17 | 0.21 | 0.30 | 0.26 | - |

Note: MA = mathematics anxiety, GA = general anxiety, Int = mathematics interest, S-Eff =mathematics self-efficacy, GCSE = mathematics GCSE exam score, UN = understanding numbers, PVT = problem verification test, NS = number sense, tw1 = twin1, tw2 = twin2.

**Table S4**. Intraclass correlations, heritability, shared and nonshared environmental estimates for all measures with 95% confidence intervals

| | rMZ | rDZ | A | C | D | E |
|---|---|---|---|---|---|---|
| MA | .43** | .09** | .37 (.29, .43) | - | - | .63 (.57, .70) |
| GA | .44** | .17** | .41 (.34, .48) | - | - | .59 (.52, .64) |
| INT | .43** | .18** | .43 (.37, .48) | - | - | .57 (.53, .62) |
| S-EFF | .59** | .25** | .58 (.52, .63) | - | - | .42 (.42, .46) |
| GCSE | .82** | .49** | .62 (.54, .71) | .19 (.11, .26) | - | .19 (.18, .20) |
| UN | .61** | .34** | .63 (.58, .68) | - | - | .36 (.33, .40) |
| PVT | .56** | .23** | .59 (.47, .64) | - | - | .41 (.38, .45) |
| NS | .33** | .19** | .36 (.29, .42) | - | - | .64 (.59, .69) |

Note: ** = p< .01; 95% confidence intervals in parentheses, A = additive genetic influences; D = non-additive genetic influences; C= shared environmental influences; E = nonshared environmental influences; MA = mathematics anxiety; GA= general anxiety; INT = interest; S-EFF = self-efficacy; GCSE = mathematics GCSE exam score; UN = understanding numbers; PVT = problem verification test; NS = number sense.

**Table S5**. Model fit indices for all univariate models and nested models

| | Baseline | Comparison | -2LL | df | AIC | *p* |
|---|---|---|---|---|---|---|
| (a) Mathematics Anxiety | | | | | | |
| 1 | Saturated | - | 8173.964 | 2919 | 2335.964 | NA |
| 2 | Saturated | ADE | 8180.603 | 2925 | 2330.603 | 0.356 |
| **3** | **ADE** | **AE** | **8191.810** | **2926** | **2339.810** | **0.028** |
| 4 | ACE | E | 8286.672 | 2927 | 2432.672 | 0.000 |
| (b) General Anxiety | | | | | | |
| 1 | Saturated | - | 8150.253 | 2919 | 2312.253 | NA |
| 2 | Saturated | ADE | 8154.761 | 2925 | 2304.761 | 0.608 |
| **3** | **ADE** | **AE** | **8155.145** | **2926** | **2303.145** | **0.535** |
| 4 | ADE | E | 8286.672 | 2927 | 2432.672 | 0.00 |
| (c) Mathematics interest | | | | | | |
| 1 | Saturated | - | 14008.597 | 5019 | 3970.60 | NA |
| 2 | Saturated | ADE | 14013.891 | 5025 | 3963.89 | 0.507 |
| **3** | **ADE** | **AE** | **14015.521** | **5026** | **3963.52** | **0.202** |
| 5 | ADE | E | 14244.217 | 5027 | 4190.22 | 0.000 |
| (d) Mathematics self-efficacy | | | | | | |
| 1 | Saturated | - | 13795.712 | 5020 | 3755.712 | NA |
| 2 | Saturated | ADE | 13796.995 | 5026 | 3744.995 | 0.973 |
| **3** | **ADE** | **AE** | **13798.807** | **5027** | **3744.807** | **0.178** |
| 4 | ADE | E | 14247.055 | 5028 | 4191.055 | 0.000 |
| (e) Mathematics GCSE grade | | | | | | |
| 1 | Saturated | - | 12219.407 | 4767 | 2685.407 | NA |
| **2** | **Saturated** | **ACE** | **12220.770** | **4773** | **2674.770** | **0.968** |
| 3 | ACE | AE | 12240.237 | 4774 | 2692.238 | 0.000 |
| 4 | ACE | CE | 12458.058 | 4774 | 2910.058 | 0.000 |

| | | | | | | |
|---|---|---|---|---|---|---|
| 5 | ACE | E | 13529.263 | 4775 | 3979.263 | 0.000 |
| (f) Understanding numbers | | | | | | |
| 1 | Saturated | - | 12161.345 | 4473 | 3215.345 | NA |
| 2 | Saturated | ACE | 12166.992 | 4479 | 3208.992 | 0.464 |
| **3** | **ACE** | **AE** | **12168.361** | **4480** | **3208.362** | **0.242** |
| 4 | ACE | CE | 12246.462 | 4480 | 3286.462 | 0.000 |
| 5 | ACE | E | 12695.163 | 4481 | 3733.163 | 0.000 |
| (g) Mathematics Problem Verification Test | | | | | | |
| 1 | Saturated | - | 12845.662 | 4677 | 3491.662 | NA |
| 2 | Saturated | ACE | 12848.623 | 4683 | 3482.623 | 0.814 |
| **3** | **ACE** | **AE** | **12848.623** | **4684** | **3480.623** | **1.000** |
| 4 | ACE | CE | 12927.622 | 4684 | 3559.622 | 0.000 |
| 5 | ACE | E | 13273.925 | 4685 | 3903.925 | 0.000 |
| (h) Number sense | | | | | | |
| 1 | Saturated | - | 13358.946 | 4761 | 3836.946 | NA |
| 2 | Saturated | ADE | 13370.762 | 4767 | 3836.762 | 0.066 |
| **3** | **ADE** | **AE** | **13370.762** | **4768** | **3834.762** | **1.000** |
| 4 | ADE | E | 13512.241 | 4769 | 3974.241 | 0.000 |

*Note*: -2LL = negative 2 times log likelihood; df = degrees of freedom; AIC = Akaike Information Criterion; **Best fitting model**

**Table S6**. Model fit indices for the two multivariate Cholesy decompositions exploring the association between MA, attitudes and performance; Model (a) includes variables entered in the following order: mathematics anxiety, interest, self-efficacy, exam score, understanding numbers, problem verification, number sense; Model (b) includes variables entered in the following order: exam score understanding numbers, problem verification, number sense, interest, self-efficacy, mathematics anxiety.

| **Models compared** | **ep** | **-2LL** | **df** | **AIC** | ***p*** |
|---|---|---|---|---|---|
| (a) | | | | | |
| Saturated - NA | 238 | 75881.583 | 31468 | 12945.58 | NA |
| **Saturated-Cholesky ACE** | **91** | **76044.425** | **31615** | **12814.43** | **0.175** |
| Cholesky ACE - Cholesky AE | 63 | 76096.793 | 31643 | 12810.79 | 0.0034 |
| (b) | | | | | |
| Saturated - NA | 238 | 75881.583 | 31468 | 12945.58 | NA |
| **Saturated - Cholesky ACE** | **91** | **76044.425** | **31615** | **12814.43** | **0.1758** |
| Cholesky ACE - Cholesky AE | 63 | 76102.906 | 31643 | 12816.91 | 0.0006 |

Note: ep = number of estimated parameters; -2LL = negative 2 times log likelihood; df = degrees of freedom; AIC = Akaike Information Criterion; **Best fitting model.**

**Table S7.** Cholesky decomposition: **standardized** genetic (A), shared environmental (C) and nonshared environmental (E) path estimates and (95% confidence intervals) for the multivariate association between mathematics anxiety, attitudes and performance

| | **A1 (95% CIs)** | **A2 (95% CIs)** | **A3 (95% CIs)** | **A4 (95% CIs)** | **A5 (95% CIs)** | **A6 (95% CIs)** | **A7 (95% CIs)** |
|---|---|---|---|---|---|---|---|
| 1. Maths Anxiety | .58 (.58; .61) | - | - | - | - | - | - |
| 2. Maths Interest | -.43 (-.46; -.42) | .47 (.47; .48) | - | - | - | - | - |
| 3. Maths Self-Efficacy | -.51 (-.55; -.46) | .18 (.18; .61) | .47 (.46; .47) | - | - | - | - |
| 4. Maths GCSE | -.56 (-.57; -.56) | .08 (.07; .18) | .31 (.24; .37) | .41 (.41; .42) | - | - | - |
| 5. Understand Numbers | -.47 (-.48; -.43) | .12 (.12; .13) | .32 (.32; .34) | .31 (.21; .32) | .25 (.15; .25) | - | - |
| 6. Maths PVT | -.52 (-.53; -.51) | .06 (.05; .18) | .22 (.22; .23) | .20 (.12; .31) | .24 (.24; .27) | .26 (.25; .26) | - |
| 7. Number Sense | -.17 (-.23; -.16) | .02 (.02; .10) | -.03 (-.03; -.02) | .17 (.04; .34) | .37 (.37; .38) | -.06 (-.07; -.05) | .33 (.33; .47) |
| | **C1 (95% CIs)** | **C2 (95% CIs)** | **C3 (95% CIs)** | **C4 (95% CIs)** | **C5 (95% CIs)** | **C6 (95% CIs)** | **C7 (95% CIs)** |
| 1. Maths Anxiety | .15 (.11; .15) | - | - | - | - | - | - |
| 2. Maths Interest | .08 (.00; .09) | .12 (.12; .13) | - | - | - | - | - |
| 3. Maths Self-Efficacy | .18 (.08; .19) | .11 (.10; .11) | .07 (.07; .09) | - | - | - | - |
| 4. Maths GCSE | .42 (.42; .43) | .18 (.18; .21) | .09 (.08; .17) | .10 (.10; .11) | - | - | - |
| 5. Understand Numbers | .34 (.31; .39) | -.06 (-.18; .20) | .07 (.05; .23) | -.02 (-.21; .23) | .00 (.00; .04) | - | - |
| 6. Maths PVT | .32 (.31; .38) | -.02 (-.04; .01) | .01 (.00; .04) | -.02 (-.04; .01) | .00 (-.02; .01) | .00 (.00; .15) | - |
| 7. Number Sense | .10 (.08; .12) | .09 (.09; .32) | .14 (-.17; .16) | -.10 (-.11; -.07) | .00 (.00; .31) | .00 (-.01; .26) | .00 (.01; .08) |
| | **E1 (95% CIs)** | **E2 (95% CIs)** | **E3 (95% CIs)** | **E4 (95% CIs)** | **E5 (95% CIs)** | **E6 (95% CIs)** | **E7 (95% CIs)** |
| 1. Maths Anxiety | .80 (.78; .80) | | | | | | |
| 2. Maths Interest | -.25 (-.25; -.24) | .72 (.70; .72) | | | | | |
| 3. Maths Self-Efficacy | -.22(-.22; -.21) | .20 (.18; .22) | .59 (.58; .60) | | | | |
| 4. Maths GCSE | -.11(-.25; -.24) | .11 (.11; .12) | .07 (.07; .08) | .39 (.38; .40) | | | |
| 5. Understand Numbers | -.10(-.14; -.07) | .10 (.07; .10) | .08 (.08; .10) | .10 (.09; .10) | .59 (.58; .60) | | |
| 6. Maths PVT | -.11(-.12; -.10) | .13 (.11; .13) | .10 (.10; .13) | .13 (.12; .15) | .09 (.08; .09) | .59 (.57; .60) | |
| 7. Number Sense | -.02(-.03; -.01) | .02 (.02; .04) | .09 (.05; .13) | .07 (.07; .12) | .05 (.05; .06) | .10 (.10; .11) | .79 (.78; .80) |

**Table S8.** Cholesky decomposition: **standardized squared** genetic (A), shared environmental (C) and nonshared environmental (E) path estimates and (95% confidence intervals) for the multivariate association between mathematics performance, attitudes and MA

| | A1 (95% CIs) | A2 (95% CIs) | A3 (95% CIs) | A4 (95% CIs) | A5 (95% CIs) | A6 (95% CIs) | A7 (95% CIs) |
|---|---|---|---|---|---|---|---|
| 1. Maths GCSE | 0.58 (0.52, 0.59) | | | | | | |
| 2. Understand Numbers | 0.42 (0.41, 0.43) | 0.07 (0.06, 0.08) | | | | | |
| 3. Maths PVT | 0.34 (0.33, 0.42) | 0.06 (0.02, 0.06) | 0.09 (0.08, 0.10) | | | | |
| 4. Number Sense | 0.04 (0.01, 0.04) | 0.10 (0.09, 0.11) | 0.00 (0.00, 0.07) | 0.16 (0.02, 0.29) | | | |
| 5. Maths Interest | 0.13 (0.12, 0.14) | 0.01 (0.00, 0.01) | 0.00 (0.00, 0.00) | 0.00 (0.00, 0.00) | 0.26 (0.06, 0.31) | | |
| 6. Maths Self-Efficacy | 0.33 (0.32, 0.34) | 0.01 (0.00, 0.05) | 0.00 (0.00, 0.00) | 0.04 (0.00, 0.05) | 0.02 (.00, .02) | 0.10 (.00, .11) | |
| 7. Maths Anxiety | 0.18 (0.17, 0.18) | 0.00 (0.00, 0.00) | 0.04 (0.00, 0.07) | 0.00 (0.00, 0.00) | 0.03 (0.00, 0.03) | 0.01 (.00, .01) | 0.08 (.00, .09) |
| | **C1 (95% CIs)** | **C2 (95% CIs)** | **C3 (95% CIs)** | **C4 (95% CIs)** | **C5 (95% CIs)** | **C6 (95% CIs)** | **C7 (95% CIs)** |
| 1. Maths GCSE | 0.22 (0.16, 0.23) | | | | | | |
| 2. Understand Numbers | 0.08 (0.06, 0.09) | 0.04 (0.03, 0.05) | | | | | |
| 3. Maths PVT | 0.07 (0.03, 0.11) | 0.03 (0.02, 0.06) | 0.00 (0.00, 0.00) | | | | |
| 4. Number Sense | 0.02 (0.00, 0.02) | 0.00 (0.00, 0.00) | 0.00 (0.00, 0.12) | 0.02 (0.00, 0.12) | | | |
| 5. Maths Interest | 0.01 (0.00, 0.05) | 0.00 (0.00, 0.00) | 0.00 (0.00, 0.00) | 0.00 (0.00, 0.06) | 0.00 (0.00, 0.00) | | |
| 6. Maths Self-Efficacy | 0.04 (0.04, 0.09) | 0.00 (0.00, 0.00) | 0.00 (0.00, 0.01) | 0.00 (0.00, 0.04) | 0.00 (0.00, 0.00) | 0.00 (0.00, 0.00) | |
| 7. Maths Anxiety | 0.02 (0.01, 0.06) | 0.00 (0.00, 0.00) | 0.00 (0.00, 0.05) | 0.00 (0.00, 0.04) | 0.00 (0.00, 0.00) | 0.00 (0.00, 0.00) | 0.00 (0.00, 0.00) |
| | **E1 (95% CIs)** | **E2 (95% CIs)** | **E3 (95% CIs)** | **E4 (95% CIs)** | **E5 (95% CIs)** | **E6 (95% CIs)** | **E7 (95% CIs)** |
| 1. Maths GCSE | 0.17 (0.16, 0.20) | | | | | | |
| 2. Understand Numbers | 0.02(0.01, 0.02) | 0.35 (0.32, 0.38) | | | | | |
| 3. Maths PVT | 0.04 (0.03, 0.05) | 0.01 (0.01, 0.02) | 0.34 (0.32,0 .38) | | | | |
| 4. Number Sense | 0.01 (0.00, 0.01) | 0.00 (0.00, 0.01) | 0.01 (0.00, 0.01) | 0.62 (.56, .69) | | | |
| 5. Maths Interest | 0.06 (0.06, 0.07) | 0.01 (0.01, 0.01) | 0.01 (0.00, 0.01) | 0.00 (0.00, 0.00) | 0.49 (0.48, 0.54) | | |
| 6. Maths Self-Efficacy | 0.04 (0.03, 0.05) | 0.01 (0.00, 0.01) | 0.01 (0.00, 0.01) | 0.00 (0.00, 0.00) | 0.03 (0.03, 0.04) | 0.34 (0.34, 0.36) | |
| 7. Maths Anxiety | 0.04 (0.03, 0.05) | 0.01 (0.00, 0.01) | 0.01 (0.00, 0.01) | 0.00 (0.00, 0.00) | 0.03 (003, 0.06) | 0.02 (0.01, 0.02) | 0.52 (0.51, 0.59) |

**Table S9**. Standardised paths for the Cholesky decomposition exploring the origins of the association between general anxiety, mathematics anxiety, mathematics attitudes and mathematics performance.

| | **A1 (95% CIs)** | **A2 (95% CIs)** | **A3 (95% CIs)** | **A4 (95% CIs)** | **A5 (95% CIs)** | **A6 (95% CIs)** | **A7 (95% CIs)** | **A8 (95%** |
|---|---|---|---|---|---|---|---|---|
| 1. General Anxiety | .58 (.57; .58) | - | - | - | - | - | - | - |
| 2. Maths Anxiety | .29 (.28; .29) | .51 (.50; .53) | - | - | - | - | - | - |
| 3. Maths Interest | -.08 (-.09; -.07) | -.45 (-.45; -.44) | .44 (.43; .44) | - | - | - | - | - |
| 4. Maths Self-Efficacy | -.17 (-.17; -.17) | -.49 (-.49; -.48) | .14 (.12; .15) | .46 (.45; .47) | - | - | - | - |
| 5. Maths GCSE | -.21 (-.22; -.16) | -.53 (-.53; -.52) | .05 (.04; .07) | .32 (.31; .32) | .41 (.41; .41) | - | - | - |
| 6. Understand Numbers | -.14 (-.14; -.12) | -.47 (-.47; -.47) | .09 (.08; .09) | .32 (.32; .32) | .29 (.28; .37) | .24 (.24; .26) | - | - |
| 7. Maths PVT | -.17 (-.18; -.17) | -.51 (-.52; -.49) | .01 (-.07; .12) | .21 (.20; .22) | .20 (.20; .20) | .24 (.23; .24) | .24 (.23; .24) | |
| 8. Number Sense | .03 (-.06; .03) | -.22 (-.22; -.15) | -.05 (-.09; -.04) | -.07 (-.08; -.01) | .19 (.18; .19) | .39 (.28; .40) | -.19 (-.19; -.18) | .00 (-.00; |
| | **C1 (95% CIs)** | **C2 (95% CIs)** | **C3 (95% CIs)** | **C4 (95% CIs)** | **C5 (95% CIs)** | **C6 (95% CIs)** | **C7 (95% CIs)** | **C8 (95%** |
| 1. General Anxiety | .25 (.25; .34) | - | - | - | - | - | - | - |
| 2. Maths Anxiety | .04 (.04; .06) | .15 (.14; .15) | - | - | - | - | - | - |
| 3. Maths Interest | -.05 (-.06; -.03) | .11 (.10; .11) | .10 (.10; .11) | - | - | - | - | - |
| 4. Maths Self-Efficacy | -.05 (-.05; -.04) | .20 (.16; .21) | .08 (.05; .10) | .07 (.00; .09) | - | - | - | - |
| 5. Maths GCSE | .12 (.12; .12) | .39 (.39; .46) | .19 (.18; .30) | .14 (-.08; .16) | .10 (.08; .11) | - | - | - |
| 6. Understand Numbers | .13 (.12; .13) | .31 (.30; .36) | -.06 (-.11; .09) | .07 (.06; .08) | .00 (.00; .04) | .00 (-.23; .05) | - | - |
| 7. Maths PVT | .06 (.06; .07) | .32 (.26; .32) | -.03 (-.03; -.01) | .07 (.06; .08) | .00 (-.18; .03) | .00 (.00; .01) | .00 (.00; .01) | |
| 8. Number Sense | -.17 (-.18; .08) | .17 (.16; .18) | .00 (-.01; .01) | .00 (-.05; .17) | .00 (-.19; .19) | .00 (-.07; .13) | .00 (-.03; .03) | .00 (.00; .23) |
| | **E1 (95% CIs)** | **E2 (95% CIs)** | **E3 (95% CIs)** | **E4 (95% CIs)** | **E5 (95% CIs)** | **E6 (95% CIs)** | **E7 (95% CIs)** | **E8 (95%** |
| 1. General Anxiety | .78 (.77; .78) | - | - | - | - | - | - | - |
| 2. Maths Anxiety | .18 (-17; .19) | .78 (.77; .78) | - | - | - | - | - | - |
| 3. Maths Interest | -.03(-.03; -.01) | -.25 (-.26; -.23) | .72 (.71; .72) | - | - | - | - | - |
| 4. Maths Self-Efficacy | -.02(-.03; -.01) | -.22 (-.25; -.22) | .20 (.20; .22) | .59 (.59; .61) | - | - | - | - |
| 5. Maths GCSE | -.01(-.01; .01) | -.11 (-.12; -.11) | .11 (.11; .13) | .07 (.07; .07) | .39 (.39; .39) | - | - | - |
| 6. Understand Numbers | -.02(-.03; .01) | -.10 (-.10; -.09) | .10 (.10; .13) | 08 (.08; .09) | .10 (.09; .10) | .59 (.58; .59) | - | - |
| 7. Maths PVT | .00 (.00; .01) | -.12 (-.12; -.11) | .13 (.12; .13) | .10 (.10; .10) | .12 (.12; .15) | .09 (.08; .09) | .59 (.58; .60) | - |
| 8. Number Sense | .00 (-.01; .01) | -.02 (-.02; .02) | .03 (.01; .04) | .10 (.10; .12) | .07 (.07; .07) | .05 (.01; .06) | .11 (.10; .12) | .79 (.79; .80) |
